# Codon-optimized FAM132b prevents diet-induced obesity by modulating adrenergic response and insulin action

**DOI:** 10.1101/2020.08.31.275503

**Authors:** Zhengtang Qi, Jie Xia, Xiangli Xue, Wenbin Liu, Zhuochun Huang, Xue Zhang, Yong Zou, Jianchao Liu, Jiatong Liu, Xingtian Li, Lu Cao, Lingxia Li, Zhiming Cui, Benlong Ji, Qiang Zhang, Shuzhe Ding, Weina Liu

## Abstract

FAM132b, also known as myonectin, has been identified as a myokine produced by exercise. It is a secreted protein precursor that belongs to the adipolin/erythroferrone family, and has hormone activity in circulation to regulate cellular iron homeostasis and lipid metabolism via unknown receptors. Here, adeno-associated viral vectors (AAV9) were engineered to induce overexpression of FAM132b with 2 codon mutations (A136T and P159A). Treatment of mice under high-fat diet feeding with FAM132b gene transfer resulted in marked reductions in body weight, fat depot, adipocytes size, glucose intolerance and insulin resistance. Moreover, FAM132b overproduction reduced glycemic response to epinephrine (EPI) in whole body and increased lipolytic response to EPI in adipose tissues. This adrenergic response of adipose tissue led to the result that gene transfer reduced glycogen utilization and increased fat consumption in skeletal muscle during exercise. FAM132b knockdown by shRNA significantly increased glycemic response to EPI in vivo and reduced adipocytes response to EPI and adipose tissue browning. Structural analysis suggested that FAM132b mutants delivered by AAV9 may form a weak bond with ADRB2, and potentially bind to insulin against insulin receptor by blocking the receptor binding sites on insulin B-chain. Our study underscores the potential of FAM132b gene therapy with codon optimization to treat obesity by modulating adrenergic response and interfering insulin action.

**Significance:** We show here that AAV9-mediated expression of FAM132b with A136T and P159A is a safe and effective therapeutic strategy for improving glucose homeostasis. This is the first demonstration of a therapeutic effect on metabolic disorders in mice with FAM132b codon optimization. These therapeutic effects indicate that FAM132b gene transfer with selective codon mutants in vivo might be a valid therapy for diabetes that can be extended to other metabolic disorders.

## Introduction

Hyperglycemia and diabetes are associated with the activation of hypothalamic-pituitary-adrenal (HPA) axis (1). Recent studies demonstrated that suppression of the HPA axis improves glycemic control and reverses diabetes (2, 3). Chronic stress increased sympathetic nervous activity and elevated basal corticosterone levels and rapid glucocorticoid production, so stress exposure increases blood glucose via the activation of HPA axis (4). However, chronic exercise enhanced HPA-axis negative feedback and thus reduced HPA responses to stress or next bout of exercise (5). Thus, chronic stress is associated with a higher risk of type 2 diabetes (6), while regular exercise is helpful for glycemic control in the previous cohort studies (7). High-fat feeding not only causes glucose intolerance, insulin resistance and metabolic disorders, and also enhances HPA axis tone in a manner that resembles HPA hyperactivity in chronic stress exposure (8, 9). Evidence from chronic stress suggests lowering glycemic response to HPA axis by some means contributes to glycemic control in patients with diabetes or hyperglycemia.

The important feature of childhood and adolescent obesity is sympathetic overactivity and in vivo resistance of lipolysis to catecholamines (10, 11). The activation in the HPA axis is intended to increase beta-adrenergic thermogenesis and expend excess energy, but adipose tissue insensitivity to adrenergic stimulation in young age leads to fat storage and even adiposity. In turn, sympathetic activity could be further enhanced as a consequence of accumulating adiposity in order to prevent further fat storage (12). In obese subjects, exercise increases sympathetic and adrenergic activity, but induces a desensitization in β1- and β2-adrenergic lipolytic pathways and the stimulation of α2-mediated antilipolytic action in adipose tissue (13, 14). This suggests that obese people exhibit lipolysis resistance to exercise by modulating the activity of adrenergic receptors (ARs). Hepatic gluconeogenesis and adipocyte lipolysis are controlled coordinately by HPA axis. The higher rate of hepatic gluconeogenesis is attributed to hypoleptinemia-induced activity of the HPA axis, resulting in higher rates of adipocyte lipolysis (2). Thus, it is of great significance to explore a novel method to improve adrenergic response of adipose tissue for improving glucose homeostasis and obesity.

Gene therapy targeting pancreatic islet and liver has emerged as one of the potential approaches to treat autoimmune diabetes and type 2 diabetes (15-17). For example, neuroD-betacellulin gene therapy induces islet neogenesis in the liver and reverses insulin-dependent diabetes in mice (18). A single injection of an adeno-associated virus (AAV) serotype 8 vector carrying human insulin gene induced hepatic production of insulin and corrected streptozotocin (STZ)-induced diabetes in mice (16). In high-fat diet (HFD) or STZ-induced model of T2DM, lentivirus-mediated glucagon-like peptide-1 gene therapy reduced blood glucose levels, along with improving insulin sensitivity and glucose tolerance (19). Additionally, AAV-mediated gene transfer of fibroblast growth factor 21 (FGF21), bone morphogenetic protein-4 (BMP4) and a glucagon-like peptide-1 receptor agonist Exendin 4 has been identified as a strategy to fight obesity and metabolic diseases (20-22). In these gene therapies, AAV was genetically manipulated and packaged to induce liver, adipose tissue, or skeletal muscle to secrete cytokines and target proteins, which have hormone activity to regulate glucose and lipid metabolism in the whole-body.

FAM132b is a secreted protein precursor that belongs to the FAM132 family. FAM132b is widely expressed in skeletal muscle, white adipose tissue (WAT), liver, hypothalamus, pituitary, adrenal gland and spleen, and weakly expressed in brown fat, heart, kidney, lung and pancreas. FAM132b is also known as erythroferrone (Erfe, produced by erythroblasts) and myonectin (produced by skeletal muscle), C1QTNF15, and CTRP15 (23-25). FAM132b has hormone activity in extracellular region. Binding of FAM132b to an unknown receptor regulates cellular iron homeostasis, fatty acid transport, fatty acid metabolic process, and inflammation. Erfe-deficient mice exhibited severe anemia, lower hemoglobin content, lower circulating iron level, higher hepcidin expression, and abnormal liver physiology (26). Myonectin (CTRP15) has been identified as a myokine that links skeletal muscle to systemic lipid homeostasis and suppresses hepatic autophagy, but higher level of circulating FAM132b predicts the development of type 2 diabetes (23). Moreover, myonectin functions as an exercise-induced myokine which ameliorates myocardial ischemic injury by suppressing apoptosis and inflammation in heart (27). These evidence corroborate that FAM132b may be an attractive candidate for gene therapy to combat obesity and T2DM.

Here, we took advantage of adeno-associated virus serotype 9 (AAV9) and its potential to delivery FAM132b in vivo to develop a gene therapy against obesity and T2DM. In the process of virus packaging, codon optimization was performed on FAM132b due to the need of vector construction, leading to mutation of A136T and P159A. We demonstrate that an administration of AAV9 encoding FAM132b with A136T and P159A enabled an increase in FAM132b levels in circulation, liver and skeletal muscle. Gene transfer resulted in improved glucose tolerance, attenuated HPA axis activity, increased glucose uptake, and reduced lipolysis resistance to epinephrine (EPI) in HFD-fed mice. However, FAM132b gene therapy attenuated glycemic responses to EPI and promoted lipolysis in response to EPI and exercise. Our results indicate that FAM132b gene transfer with codon mutations improves adrenergic responses of whole body and thus prevents obesity and metabolic disorders.

## Results

### In vivo gene transfer by rAAV9 elevates FAM132b levels in circulation, liver and skeletal muscle

To study the impact of FAM132b overexpression, the codon-optimized DNA sequence encoding full-length FAM132b gene was packaged into an AAV9 vector (AAV:132b; Fig. 1A). To confirm proper transgene expression by AAV-132b vector in vitro, HEK293 cells were infected with viral vectors, and the target protein was detected in cell lysates by western blot (Fig. 1B). FAM132b could also be detected by ELISA in both cell lysate and media, confirming that immunoreactive FAM132b was expressed and secreted (Fig. 1C).

**Figure 1.**
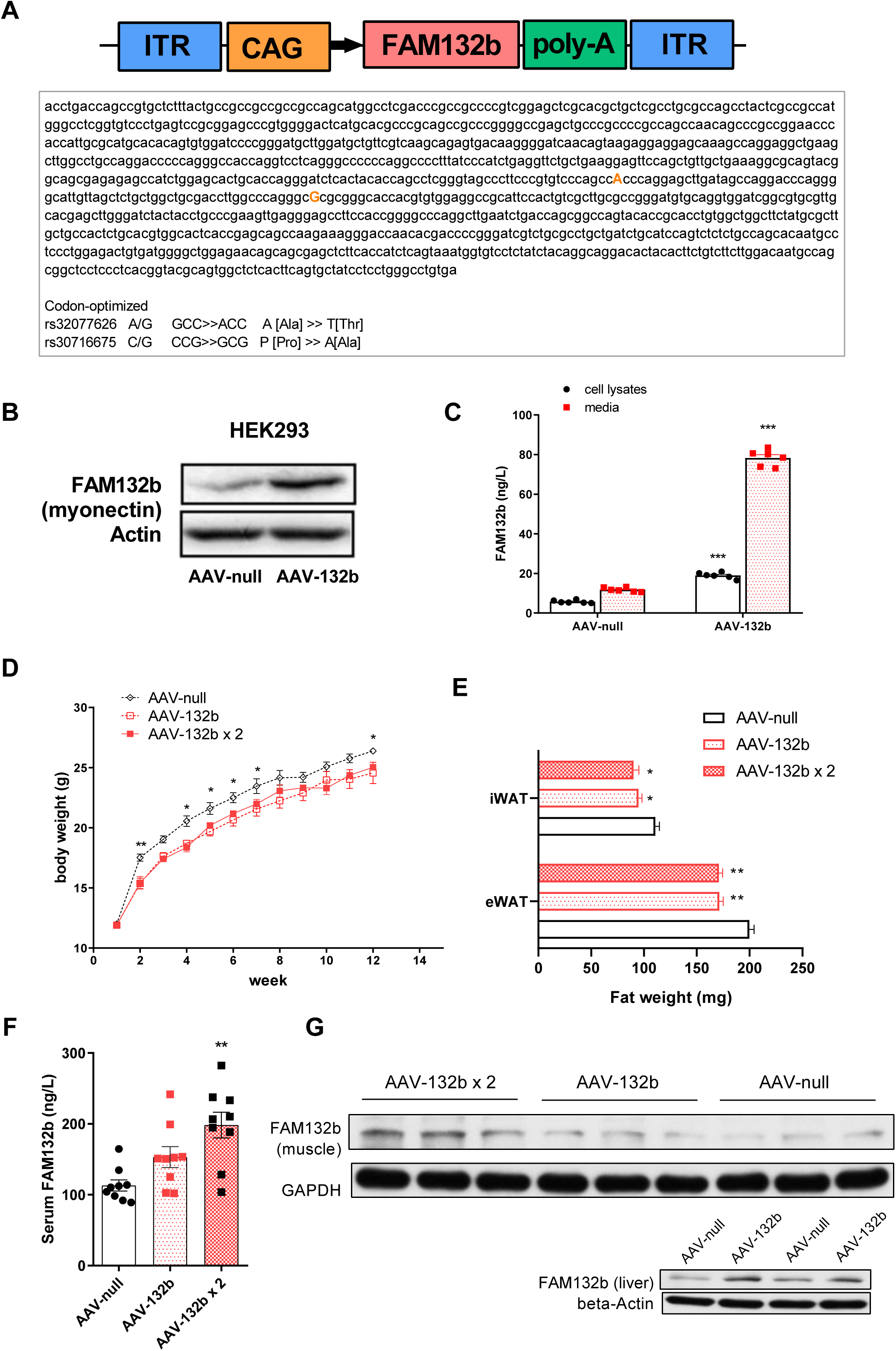
AAV9-mediated gene transfer of FAM132b counteracts HFD-induced obesity. (A) cDNA of mouse FAM132b was subcloned into EcoRI and XbaI restriction sites resulting in a 4654 bp fragment between 5′-ITR and 3′-ITR. The restriction enzymes are used for investigation of the accuracy of FAM132b insertion into constructed plasmids. ITR sequence is necessary for packaging of the transgene into virions and its integration into the host genome. (B) Western blot of FAM132b in transfected HEK293 cells with constructed plasmids. (C) Transfected HEK293 cells showed higher FAM132b levels in cell lysates and media. (D) Evolution of body weight in animals treated with once or twice infusion of AAV-132b vectors. n = 10–12/group. (E) Weight of the epididymal (eWAT) and inguinal (iWAT) white adipose tissue depots obtained from mice treated with AAV-132b vectors. n = 10–12/group. (F) Circulating levels of FAM132b after vector administration. (G) Western blot of FAM132b in transfected liver and gastrocnemius muscle after vector administration. All values are expressed as mean ± SEM. In Fig.1C, data were analyzed by two-way ANOVA with Tukey’s post hoc correction. ***P < 0.001 versus AAV-null. In Fig.1D-F, data were analyzed by one-way ANOVA with Tukey’s post hoc correction. *P < 0.05, **P < 0.01 versus AAV-null.

For in vivo experiments, 4-week-old C57BL/6J mice were fed HFD and injected 4.7×10^10^ vector or double per mouse with AAV-132b or AAV-null vehicle by the tail vein. Mice were followed closely after a single vector administration to assess effect and identify toxicity. 12 weeks after AAV9 administration, mice treated with AAV-132b had a lower body weight gain (Fig.1D), lower fat mass (Fig.1E), and higher FAM132b levels in circulation than null-treated mice (Fig.1F). Western blot analysis using anti-FAM132b antibody (AVISCERA BIOSCIENCE, A003938-03-100, Santa Clara, CA, USA) revealed FAM132b overexpression in gastrocnemius muscle and liver of mice treated with AAV-132b (Fig.1G). Double dose of gene transfer increased FAM132b expression in skeletal muscle more significantly than a single dose (Fig.1G). These experiments showed that the infusion of AAV-132b by tail vein resulted in a successful gene transfer into skeletal muscle and liver.

### FAM132b gene therapy reduces glycemic response to EPI and suppresses HPA axis activity

To understand the longitudinal change in glucose homeostasis in the development of diet-induced obesity, we performed glucose, insulin and EPI tolerance test in HFD-fed mice every week after diet intervention. At the 2^nd^ week, glucose intolerance emerged firstly in HFD-fed mice, but the difference in insulin and EPI sensitivity was not found in insulin and EPI tolerance test (Fig. S1A). Such results reappeared in the 3^rd^ week (Fig. S1B). At the 4^th^ week, glucose intolerance was further aggravated; moreover, HFD had led to insulin resistance (Fig. S1C). Blood glucose increased within 30 min after EPI injection and then began to fall back in normal chow-fed mice, but blood glucose increased for 60 min before it began to fall back in HFD-fed mice (Fig. S1C). The peak blood glucose of mice on high-fat diet was significantly higher than that of mice on normal diet (Fig. S1C), suggesting that HFD increased the glycemic response to EPI. The area under the curve (AUC) also showed that a high-fat diet caused glucose intolerance first, followed by insulin resistance and epinephrine hypersensitivity (Fig. S1D). Our data suggest that elevated EPI sensitivity, along with reduced insulin sensitivity, may further aggravate glucose intolerance in mice exposed to HFD.

Two weeks after the AAV9 injection, we evaluated glucose homeostasis in mice challenged by intraperitoneal injection of glucose, insulin, and EPI. Compared with AAV-null vehicle control group, FAM132b gene therapy significantly improved glucose intolerance and insulin resistance (Fig. 2A, 2B). FAM132b gene therapy reversed the hyperglycemic response to EPI. Within 30 minutes after EPI injection, blood glucose levels in null-treated mice surged to nearly 20mmol/L, but FAM132b gene therapy attenuated this response (Fig. 2C). Leptin is a cytokine secreted by adipose tissue and associated with obesity and metabolic disorders. Also, FAM132b gene transfer reduced serum levels of leptin and triglyceride (Fig. 2D). However, there were no significant changes in serum glucose, insulin and glycerol levels (Fig. 2D). High-fat diet was reported to enhance HPA axis activity (8, 9), leading to hyperglycemic response to EPI (28). In this study, we further found that FAM132b gene therapy reduced serum levels of corticosterone, ACTH, norepinephrine (NE), and EPI (Fig. 2E). Parameters of HPA functionality at the level of mRNA transcripts were reduced in hypothalamus of mice treated with AAV-132b (Fig. 2F), implying an inhibition of HPA axis. These results indicated that FAM132b gene therapy improved glucose homeostasis and reduced hyperglycemic response by suppressing HPA axis activity in mice fed a high-fat diet.

**Figure 2.**
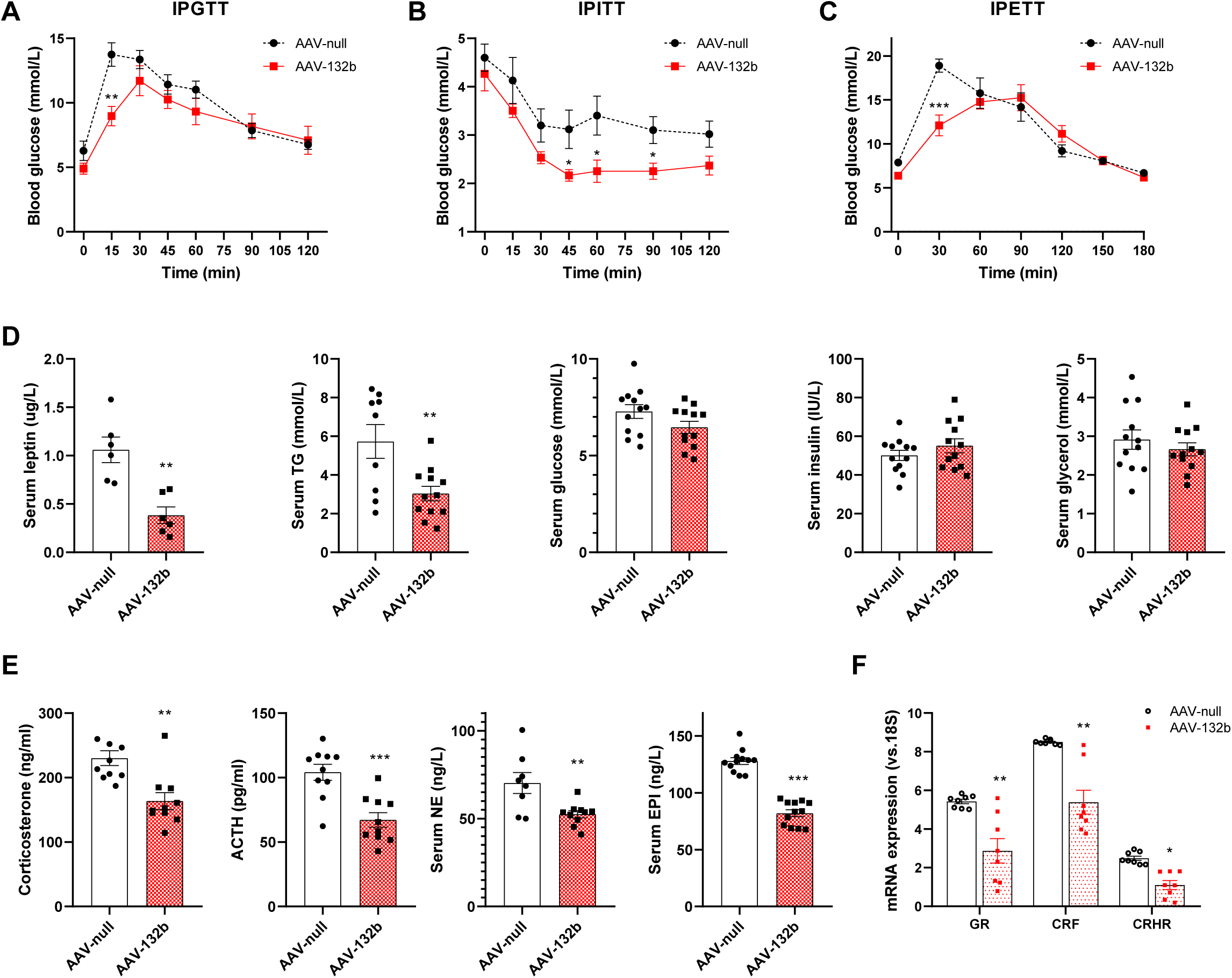
FAM132b gene transfer improves glucose homeostasis and attenuates adrenergic response of blood glucose. (A) Glucose tolerance was studied in the group of mice that initiated the HFD feeding and received once FAM132b vectors after an intraperitoneal injection of glucose (1 g/kg body weight, n = 7/group). (B) Insulin sensitivity was determined in all experimental groups after an intraperitoneal injection of insulin (0.75 units/kg body weight, n = 7/group). (C) Epinephrine sensitivity was determined by blood glucose in all experimental groups after an intraperitoneal injection of epinephrine (0.1 mg/kg body weight, n = 7/group). (D) Fasted serum levels in leptin, TG, glucose, insulin and glycerol in mice fed HFD and injected with either AAV-null or AAV-132b vectors. (E) Fasted serum levels in corticosterone, ACTH, NE, and EPI in the same groups as in (D) (F) Quantification by qRT–PCR of the expression of GR, CRF and CRHR in the hypothalamus in the group of animals that initiated the HFD feeding and received once FAM132b vectors. All values are expressed as mean ± SEM. In Fig.2A-C, data were analyzed by two-way ANOVA with Tukey’s post hoc correction. In Fig.2D-F, data were analyzed by student’s T test. *P < 0.05, **P < 0.01, ***P < 0.001 versus AAV-null group.

### FAM132b gene therapy augments insulin-stimulated glucose uptake in adipose tissue

To further explore the mechanism of improved glucose tolerance in mice treated with AAV9, we measured insulin-stimulated glucose uptake of skeletal muscle, liver, eWAT and BAT in vitro. In these tissues incubated with insulin, FAM132b gene therapy enhanced insulin-stimulated glucose uptake in the eWAT, but not in the liver and skeletal muscle (Fig. 3A). In this experiment, we used a normal diet as a control. It is noteworthy that HFD inhibited the glucose uptake of BAT, but FAM132b gene therapy reversed this effect (Fig. 3A). Western blot analysis indicated that FAM132b gene therapy increased Akt phosphorylation at Ser473 in the eWAT and BAT (Fig. 3B, 3C), suggesting the activation of insulin signaling in adipose tissues. This was consistent with the insulin-stimulated glucose uptake in the eWAT and BAT (Fig. 3A). However, these results were not observed in skeletal muscle (Fig. 3B, 3C). Furthermore, we found that Akt phosphorylation in skeletal muscle was at higher level in all groups and not significantly affected by gene transfer (Fig. 3B, 3C). In contrast, FAM132b gene therapy had a greater effect on insulin signaling in adipose tissue than in skeletal muscle. This can lead to a shift in fuel selection in adipose tissue. Thus, FAM132b augments insulin signaling in adipose tissue to increase glucose uptake, and thus improve glucose homeostasis of mice on HFD.

**Figure 3.**
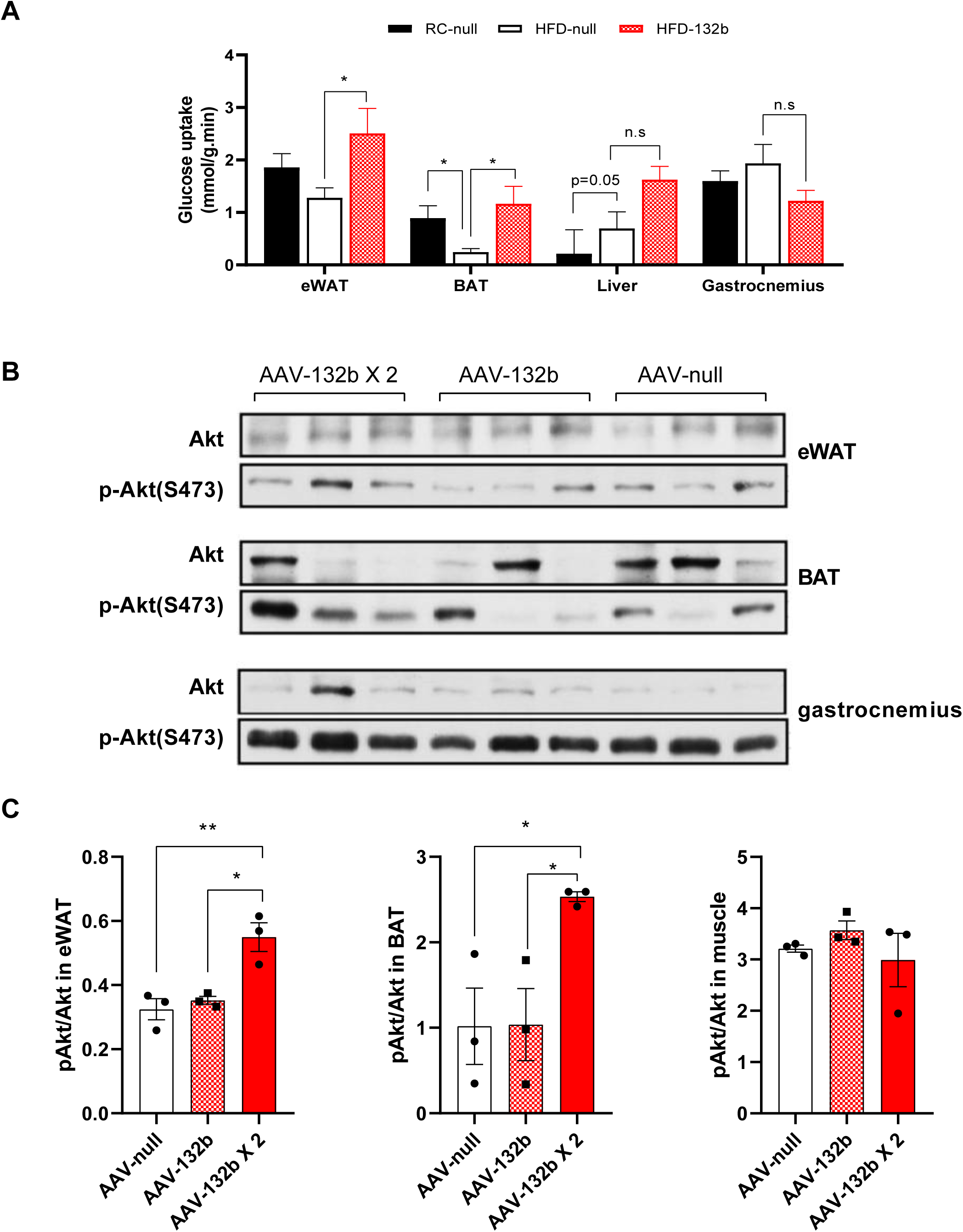
FAM132b gene transfer increases insulin-stimulated glucose uptake in eWAT and BAT. (A) Insulin-stimulated glucose uptake in eWAT, BAT, liver and gastrocnemius was evaluated in mice fed regular chow, HFD and injected with either AAV-null or AAV-132b vectors (n = 6/group). (B) Western blot analysis of Akt phosphorylation in eWAT, BAT, and gastrocnemius muscle after once or twice vector administration (n = 3/group). (C) The histogram depicts the densitometric analysis of immunoblots as shown in (B). All values are expressed as mean ± SEM. In Fig.3A, C, data were analyzed by one-way ANOVA with Tukey’s post hoc correction. *P < 0.05, **P < 0.01 indicates the difference between two groups. n.s, no significance.

### FAM132b gene therapy reduces lipolytic resistance to EPI in adipose tissue

Obese subjects exhibit fat self-protection and lipolytic resistance during exercise (13, 14). Serum glycerol level is an important marker of fat mobilization and lipolysis. Our results showed that HFD-fed mice were more sensitive to EPI than chow-fed mice. Serum glycerol in HFD-fed mice continued to increase in 90 minutes after EPI injection, but serum glycerol level of chow-fed mice was almost unchanged (Fig. S2A). The blood glucose and serum glycerol response to EPI further suggests that high-fat diet leads to increased EPI sensitivity in whole-body. However, this effect varies from liver to adipose tissue. In this study, EPI-stimulated glycerol release in culture medium was used to evaluate lipolysis ex vivo. In the time-dependent test, we found that high-fat diet led to an increase in glycerol release from liver (Fig. S2B) and a decrease in glycerol release from eWAT (Fig. S2C) 120min after EPI was added to the medium. In the experiment where eWAT and BAT were incubated with EPI for 180min, high-fat diet reduced EPI-stimulated glycerol release (Fig. S2D). No significant changes in lactic acid release were found in skeletal muscle stimulated by EPI at different concentrations (Fig. S2E). Our data suggest that early exposure to high-fat diet leads to reduced lipolysis in adipose tissues, increased lipolysis in liver and unaltered glycogenlysis in skeletal muscle. These responses to EPI are related to adrenergic receptors in different tissues (29, 30). Reduced expression of adrenergic receptors in BAT may contribute to lipolysis resistance in mice exposed to high-fat diet (Fig. S2F).

Four weeks after the AAV9 injection, we evaluated EPI-stimulated lipolysis in mice. Compared with null-treated mice, FAM132b gene therapy increased EPI-stimulated lipolysis in whole-body at the dose of 0.1mg/kg EPI, as evidenced by increased serum glycerol 30min after EPI injection (Fig. 4A). However, there was no significant difference in serum glycerol levels at 60 and 90 min after EPI stimulation (Fig. 4A). When EPI was increased to 0.5 mg/kg, there was no significant difference in serum glycerol levels at any time (Fig. 4B). Time-dependent experiments showed that lipolytic response to EPI in the whole-body was promoted by gene therapy, but this effect was not significant at higher dose of EPI. Higher concentration of EPI may result in activation of alpha2-adrenergic receptors, which blunts epinephrine-induced lipolysis in subcutaneous adipose tissue (31). Our findings suggest that FAM132b gene therapy facilitates rapid mobilization of fat at lower dose of EPI. In the cultured adipose tissues, EPI resulted in a significant increase in NEFA release (p < 0.01 for variation with time; Fig. 4C). Also, EPI resulted in a dose-dependent increase in NEFA release (p < 0.01 for variation with EPI concentration; Fig. 4D). Considering the dynamics of the lipolytic response to EPI between groups, FAM132b gene therapy promoted adipocytes lipolysis (Fig. 4C, 4D). Moreover, these site-specific differences in lipolytic response were also shown in glycerol release (Fig. 4E, 4F). Specifically, FAM132b gene therapy increased glycerol release from eWAT at the same time point or at the same concentration of EPI (Fig. 4E, 4F). Our results suggest that FAM132b delivery reduces the lipolytic resistance to EPI in adipose tissues.

**Figure 4.**
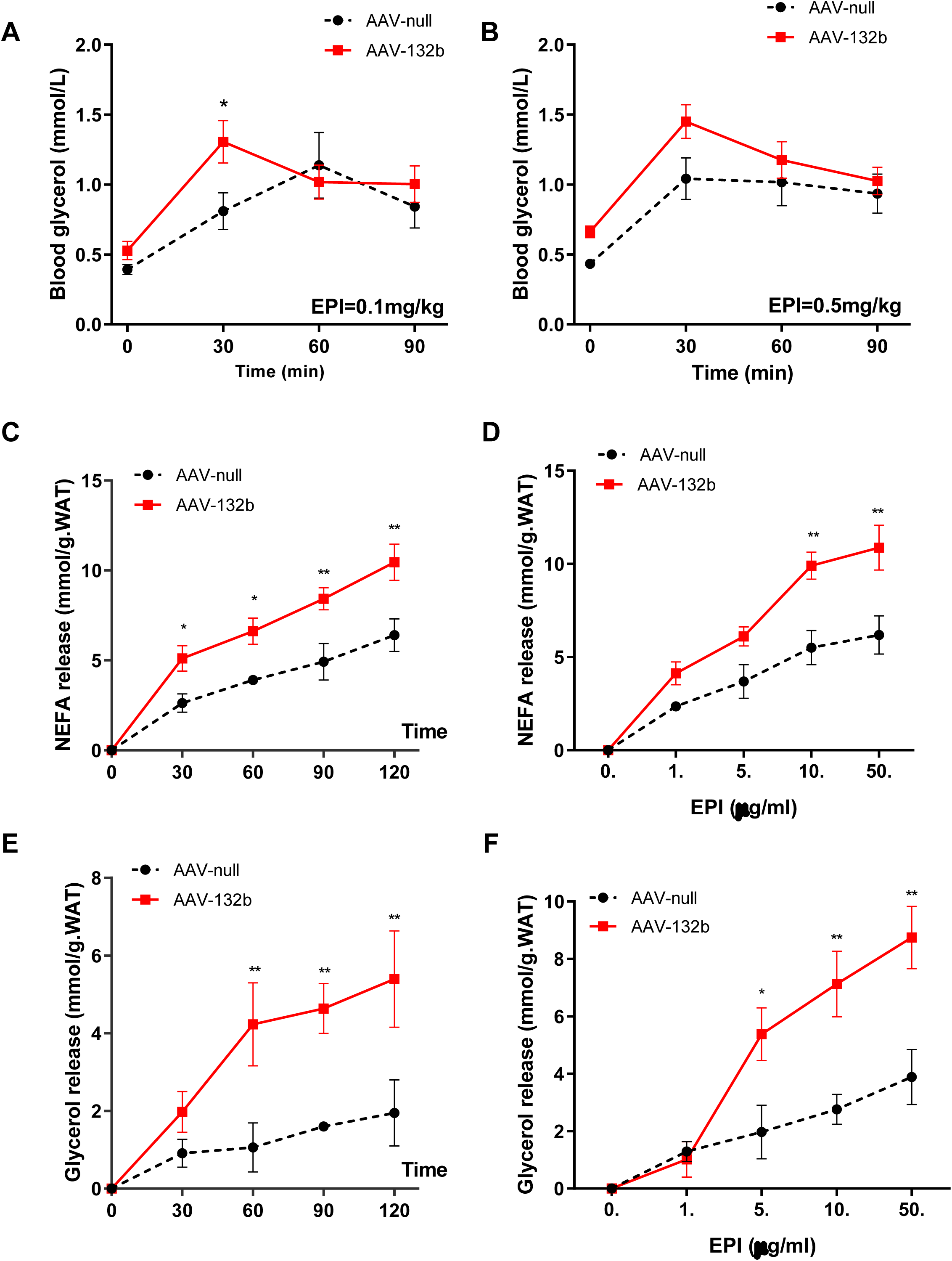
FAM132b gene transfer increases adrenergic response of lipolysis in WAT. (A) Adrenergic response of lipolysis in whole-body was determined by time-dependent blood glycerol in mice after an intraperitoneal injection of epinephrine (0.1 mg/kg body weight, n = 7/group). Mice initiated the HFD feeding and received once FAM132b vectors. (B) Adrenergic response of lipolysis in whole-body was determined by time-dependent blood glycerol in the same group of mice after an intraperitoneal injection of epinephrine (0.5 mg/kg body weight, n = 7/group). (C) Adrenergic response of lipolysis in WAT was determined by time-dependent NEFA release after EPI was added into the medium by 10 ug/ml (n = 6/group). (D) Adrenergic response of lipolysis in WAT was determined by dose-dependent NEFA release after EPI was added into the medium by the indicated concentrations (n = 6/group). (E) Adrenergic response of lipolysis in WAT was determined by time-dependent glycerol release after EPI was added into the medium by 10 ug/ml (n = 6/group). (F) Adrenergic response of lipolysis in WAT was determined by dose-dependent glycerol release after EPI was added into the medium by the indicated concentrations (n = 6/group). All values are expressed as mean ± SEM. In Fig.4A-F, data were analyzed by two-way ANOVA with Tukey’s post hoc correction. *P < 0.05, **P < 0.01 versus AAV-null group.

### FAM132b gene therapy stimulates WAT Browning

We next sought to investigate whether the increased lipolysis could be due to an increased WAT browning. FAM132b delivery with double injection (AAV-132b X2) increased emergence of adipocytes containing multilocular lipid droplets and reduced adipocyte size in the iWAT (Fig. 5A, 5C). Also, FAM132b gene transfer reduced larger lipid droplets and increased adipocyte count in the BAT (Fig. 5A, 5B). Histological analysis showed that FAM132b reduced fat storage and increased fat consumption, and double dose of AAV9:132b was more effective than a single administration. We did observe increased protein expression of UCP1, a marker for adipocytes browning, in the eWAT (Fig. 5D, 5E). In a recent study, PTEN was reported to regulate adipose tissue homeostasis and redistribution via a PTEN-leptin-sympathetic loop. Upregulation of PTEN level was associated with higher lipolysis, white fat browning, and the compensatory reduced fat mass to maintain a set point of whole-body adiposity (32). Consistent with this study, HFD-fed mice injected with AAV9:132b showed augmented WAT browning and reduced fat pad weight (Fig. 1D, 1E), as evidenced by increased PTEN levels (Fig. 5D, 5E). Increased WAT browning was further supported by elevated mRNA levels involved in lipolysis and thermogenesis. Beige/brown adipocyte markers PGC-1a, Ucp1, Vdr, Prdm16 and hormone-sensitive lipase HSL and ATGL were upregulated in the eWAT and BAT (Fig. 5F).

**Figure 5.**
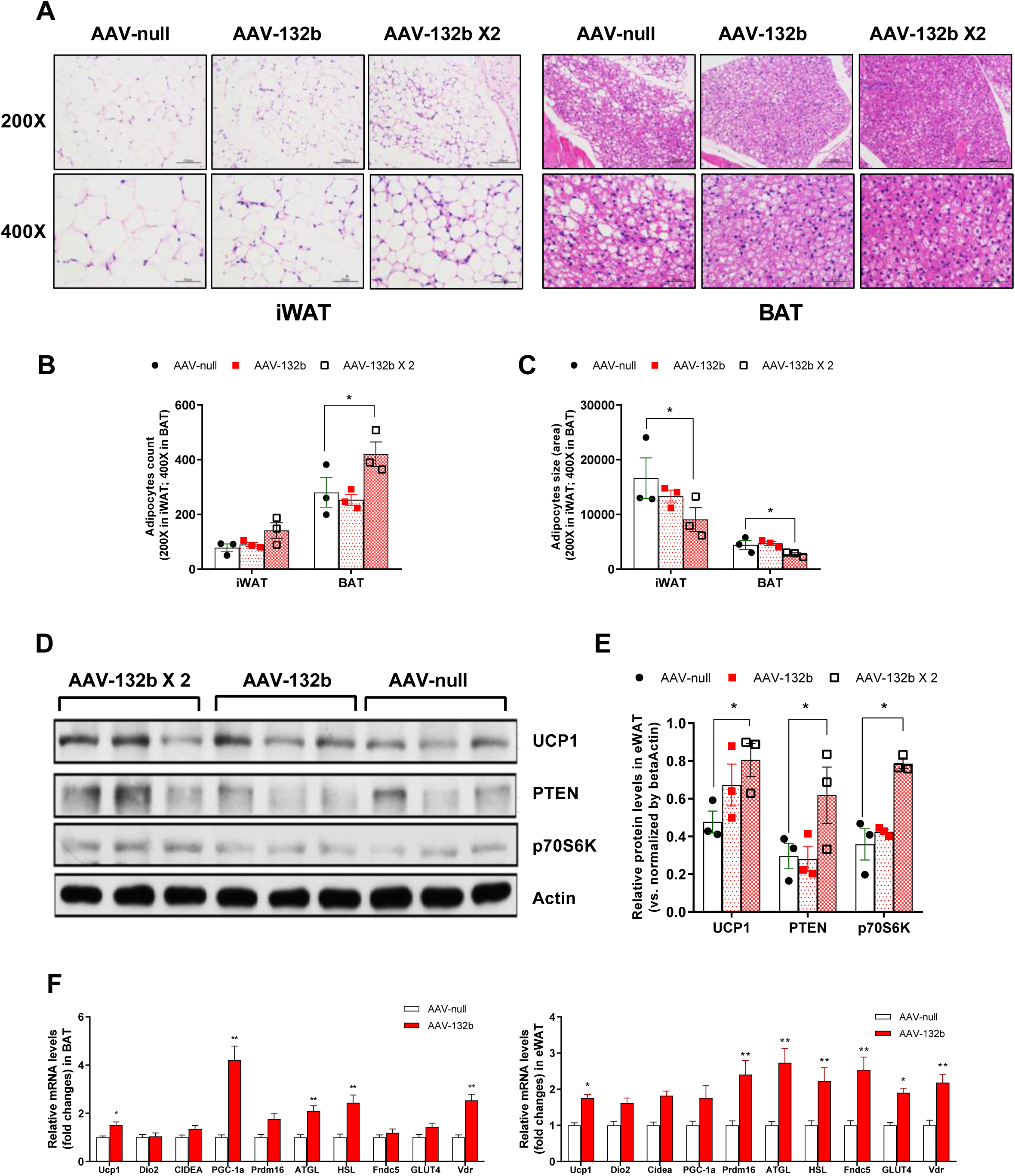
FAM132b gene transfer stimulates WAT browning. (A) Representative images of the hematoxylin-eosin staining of the iWAT (left panels) and BAT (right panels) from animals fed HFD and administered with either AAV-null or AAV-132b vectors. While HFD-fed null-injected mice had larger adipocytes, HFD-fed FAM132b-treated animals had adipocytes of reduced size. Scale bars: 100µm in 200X, 50µm in 400X. (B) Morphometric analysis of the count of adipocytes in the iWAT (200X) and BAT (400X). (C) Morphometric analysis of the size of adipocytes in the iWAT (200X) and BAT (400X). (D) Western blot analysis of UCP1, PTEN, and p70S6K in the eWAT after once or twice vector administration (n = 3/group). (E) The histogram depicts the densitometric analysis of immunoblots as shown in (D). (F) Gene expression of beige/brown adipocyte markers in the eWAT and BAT. Ucp1, uncoupling protein 1; Dio2, Deiodinase Type 2; Cidea, cell death-inducing DNA fragmentation factor-alpha-like effector A; PGC-1a, Peroxisome proliferator-activated receptor gamma coactivator-1 alpha; Prdm16, PR domain containing 16; ATGL, adipose triglyceride lipase; HSL, hormone-sensitive lipase; Fndc5, fibronectin type III domain containing 5; Glut4, glucose transporter 4; Vdr, vitamin D receptor. n = 8/group. All values are expressed as mean ± SEM. In Fig.5B, C, E, data were analyzed by one-way ANOVA with Tukey’s post hoc correction. *P < 0.05, **P < 0.01 indicates the difference between two groups. In Fig.5F, data were analyzed by student’s T test. *P < 0.05, **P < 0.01 versus AAV-null group.

### FAM132b gene therapy alters skeletal muscle fuel selection during exercise

The initial hypothesis of this study was to investigate whether gene transfer of FAM132b using AAV9 regulates skeletal muscle metabolism. However, we did not observe significant differences in insulin-stimulated glucose uptake or insulin signaling in muscles. We next evaluated the fuel selection of skeletal muscle during 40-min treadmill exercise. At rest, FAM132b gene therapy did not alter the levels of plasma glucose, NEFA, lactate, liver glycogen, muscle glycogen and malate (Fig. 6A, 6B). With 40 minutes running, mice injected with AAV9:132b had lower blood glucose and lactate, higher plasma NEFA, higher muscle and liver glycogen than AAV null-treated mice (Fig. 6A, 6B). In AAV null-treated mice, exercise elevated plasma glucose and lactate, and reduced muscle and liver glycogen; however, these changes were not significant in mice injected with AAV9:132b (Fig. 6A, 6B). This suggests that FAM132b gene therapy delayed glycogen mobilization during exercise. At rest, mice injected with AAV9:132b did not show differential metabolites of acetylcarnitine in skeletal muscle compared with AAV null-treated mice (Fig. 6C). With exercise, mice injected with AAV9:132b had higher short-chain (C:2-6), medium-chain (C:12) and long-chain acetylcarnitine (C:14-20), indicating an increase in muscle utilization of fatty acids (Fig. 6C). In addition, FAM gene therapy reduced p70S6K protein levels in skeletal muscle after one or two injections of AAV9. A single administration of AAV9 reduced PI3K protein levels, while double administration of AAV9 increased PI3K protein levels in skeletal muscle (Fig. 6D). These results in insulin signaling further suggest that FAM132 gene transfer makes skeletal muscles more prone to burn fat during exercise.

**Figure 6.**
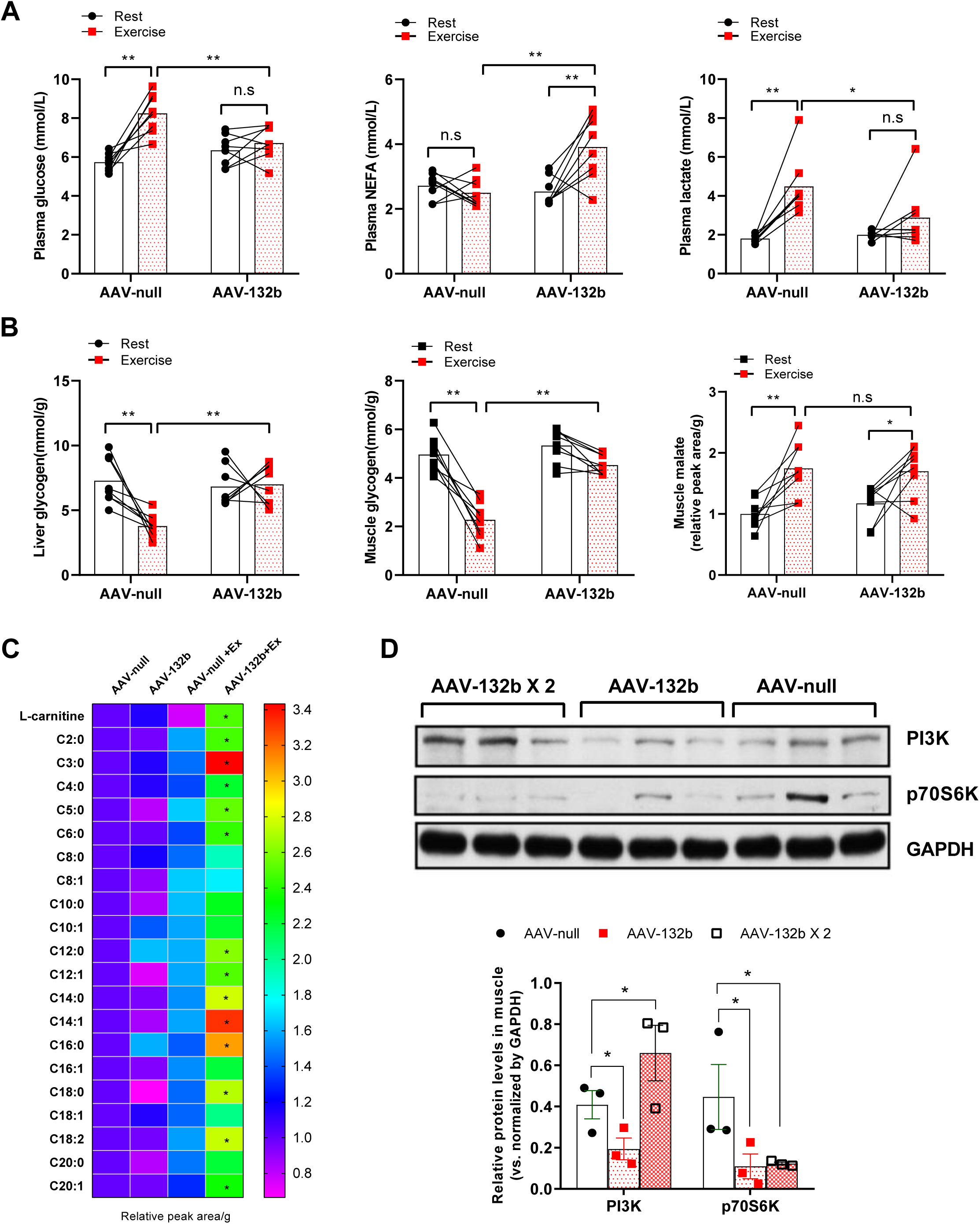
FAM132b gene transfer reduces glycogen utilization and increases fat consumption during exercise. (A) Plasma glucose, NEFA and lactate were determined during a 40-min treadmill exercise test for AAV null- and FAM132b-treated mice. n = 6/group. (B) Liver glycogen, muscle glycogen and malate were determined during a 40-min treadmill exercise test for AAV null- and FAM132b-treated mice. n = 6/group. (C) Long-chain acyl-carnitines in gastrocnemius muscle increase more with exercise in FAM132b-injected mice. Metabolite values are expressed relative to AAV-null rest for each group (n = 6). Acylcarnitines are listed by carbon chain-length. (D) Western blot analysis of PI3K and p70S6K in gastrocnemius muscle after once or twice vector administration (n = 3/group). The histogram depicts the densitometric analysis of immunoblots. All values are expressed as mean ± SEM. In Fig.6A, 6B, data were analyzed by two-way ANOVA with Tukey’s post hoc correction. *P < 0.05, **P < 0.01 indicates the difference between two groups. In Fig.6C, data were analyzed by two-way ANOVA with Tukey’s post hoc correction. * p< 0.05 between AAV-null and AAV-132b after exercise; # p<0.05 difference from baseline. In Fig.6D, data were analyzed by one-way ANOVA with Tukey’s post hoc correction. *P < 0.05, **P < 0.01 indicates the difference between two groups.

### FAM132b knockdown increases glycemic response to EPI and reduces adrenergic response of lipolysis

We further designed AAV9 with FAM132b knockdown and transfected mice by tail vein injection. As expected, AAV9 shRNA gene therapy increased the sensitivity of blood glucose to EPI. Mice fed HFD had higher blood glucose in response to EPI after AAV9 shRNA gene therapy (Fig. 7A). In contrast, AAV9 shRNA reduced glycemic response to glucose challenge in IPGTT, indicating that glucose tolerance was also improved (Fig. 7A). In EPI-stimulated adipose tissues, AAV9 shRNA reduced glycerol release from BAT (Fig. 7B). However, EPI-induced lipolysis in eWAT was not altered in mice receiving AAV9 shRNA (Fig. 7B). AAV9 shRNA did not alter insulin-stimulated glucose uptake in liver, skeletal muscle and eWAT. However, AAV9 shRNA enhanced glucose uptake in BAT (Fig. 7C). FAM132b knockdown increased serum insulin levels, but decreased serum triglyceride, EPI, NE levels (Fig. 7D). Thus, reducing FAM132b expression enhances the glycemic response to EPI, which predisposes mice to the risk of hyperglycemia during adrenergic stress. In contrast to FAM132b overexpression, FAM132b knockdown reduced thermogenic gene expression in BAT and eWAT (Fig. 7E). These data suggest that silencing FAM132b increases glycemic response to EPI in whole-body and impairs adrenergic response of lipolysis in brown adipose tissue.

**Figure 7.**
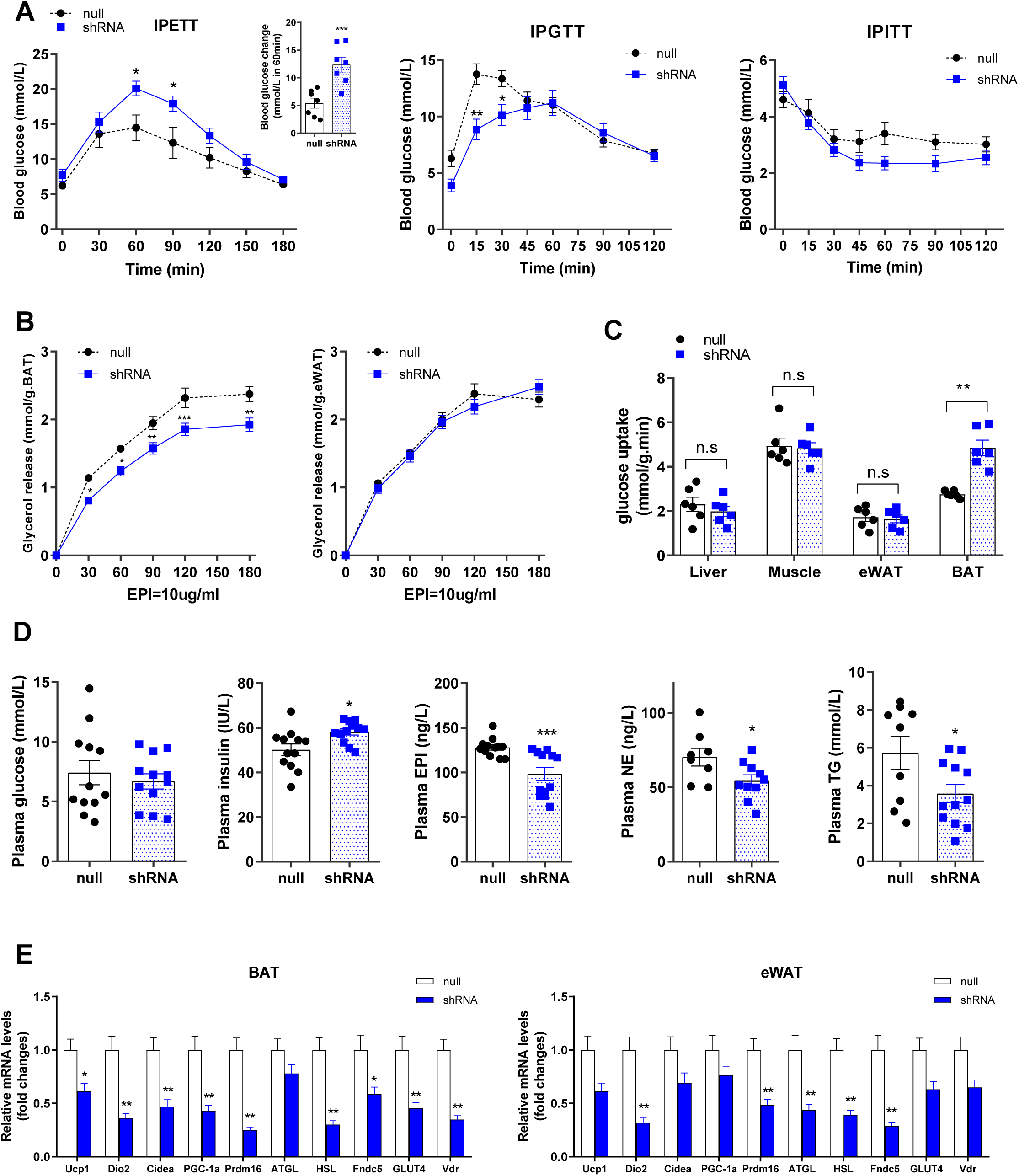
FAM132b knockdown increases adrenergic response of blood glucose and suppresses adrenergic response of lipolysis in BAT. (A) Epinephrine sensitivity, glucose tolerance, and insulin sensitivity were determined by blood glucose in 3 tolerance tests, where the group of mice that initiated the HFD feeding and received FAM132b shRNA were subjected to an intraperitoneal injection of epinephrine (0.1 mg/kg body weight), glucose (1 g/kg body weight), or insulin (0.75 units/kg body weight). n = 7/group. (B) Adrenergic response of lipolysis in BAT and eWAT was determined by time-dependent glycerol release in 180 min after EPI was added into the medium by 10 ug/ml (n = 6/group). (C) Insulin-stimulated glucose uptake in liver, gastrocnemius, eWAT and BAT was evaluated in mice fed HFD and injected with either AAV-null or AAV-132b shRNA (n = 6/group). (D) Fasted plasma levels in glucose, insulin, EPI, NE and TG in mice fed HFD and injected with either AAV-null or AAV-132b shRNA. (E) Gene expression of beige/brown adipocyte markers in the eWAT and BAT. n = 8/group. All values are expressed as mean ± SEM. In Fig.7A, 7B, data were analyzed by two-way ANOVA with Tukey’s post hoc correction. *P < 0.05, **P < 0.01, ***P < 0.001 versus AAV-null group. In Fig.7C-E, data were analyzed by student’s T test. *P < 0.05, **P < 0.01, ***P < 0.001 versus AAV-null group.

### Structural basis of codon-optimized FAM132b for the cooperative allosteric regulation of ADRB2 and insulin receptor

FAM132b has hormone activity and promotes lipid uptake into adipocytes and hepatocytes via transcriptional up-regulation of genes involved in fatty acid uptake (23). However, previous studies have not identified the receptor of FAM132b and the signal pathway to promote fatty acid transport. Our findings suggested that codon-optimized FAM132b (coFAM132b) altered the adrenergic response of blood glucose and lipolysis. We next aimed to predict the potential receptor of FAM132b and further analyze the impact of codon mutations on protein-protein interactions (PPI). Predicted secondary structure showed that random coil was dominant, and alpha helix and extended strand were less in mouse FAM132b protein (Table S2). Random coil is dispersed at N terminal: 2∼43, 54∼92, 112∼134, 139∼145, 158∼163, 187∼197, 209∼ 216, 229∼243. Codon mutations changed the secondary structure of FAM132b at 70∼72, 134∼139, 198∼201, 218∼220, 243∼254 (Table S1). Sequence 185∼340 is the subsection of the family and domains, which is defined as a specific combination of secondary structures organized into a characteristic three-dimensional structure or fold. This structure determines that FAM132b belongs to the adipolin/erythroferrone family (26). Thus, codon mutations might change the ligand properties of FAM132b.

We next modeled the full-length structure of mouse FAM132b by I-TASSER and validated its quality using PROCHECK and subjected it to energy minimization (Fig. S3). 3D structure of several potential receptors was downloaded from the protein data bank. ZDOCK was used to evaluate interactions between the native and mutant (A136T, P159A) FAM132b with these receptors. PDBePISA analysis showed that no chain contacts were found in the complexes by the docking of FAM132b with alpha-2A adrenergic receptor (PDBID: 1HLL and 6KUX), beta3 adrenergic receptor (SWISS model), GPR40 (PDBID: 5TZR). However, the docking of FAM132b and beta2 adrenergic receptor (PDBID: 3KJ6) can form a weak bond complex, and either A136T or P159A mutant will result in no chain contacts in the complex. Both mutants restore the weak bond complex with 24 hydrogen bonds and 7 salt bridges (Fig. S4A). ΔG<0 suggests that this complex may form instantaneously in vivo and function for a short time. Indeed, small-molecule drugs targeting this diverse surface have been reported to function as allosteric modulators with subtype selectivity of ADRB2 (33). Thus, the weak bond complex of FAM132b-ADRB2 impacts the binding of EPI to ADRB2 and exhibits different efficacies towards G-protein activation.

Insulin receptors on the cell surface bind to insulin to form complexes, which is the first step to increase glucose uptake and fatty acid transport via insulin signaling. Structural analysis indicates that protective hinge in insulin opens to enable its receptor engagement at B20-B23 β-turn, coupling reorientation of Phe(B24) to a 60° rotation of the B25-B28 β-strand away from the hormone core to lie antiparallel to the receptor’s L1-β2 sheet. A switch between free and receptor-bound conformations of insulin depends on the opening of this hinge, which enables conserved side chains to engage insulin receptor (34). To better address the impact of codon mutations on insulin-receptor interactions, we used ZDOCK to mock the structure of the FAM132b-insulin-receptor complex (PDBID: 4OGA). FAM132b-insulin docking showed that the family domain of FAM132b (227∼312) bound to insulin in separate spatial positions, including 40 hydrogen bonds and 10 salt bridges (11 hydrogen bonds and 5 salt bridges on the A-chain, 3 hydrogen bonds on the B-chain). In the codon mutant model, interfacing residues have 30 hydrogen bonds and 7 salt bridges (only 3 hydrogen bonds and 2 salt bridges on the B-chain, Fig. S4B). The atoms, residues, ΔG, SASA at the interface between coFAM132b and insulin were near to those at the interface between insulin and its receptor (Fig. S4B left). These results suggest that coFAM132b in circulation may compete with insulin receptors to bind insulin and thus influence insulin action.

The extent of interaction between FAM132b and insulin-receptor complex was shown in Fig. S4C. Protein-protein docking showed that the atoms, residues and solvent accessibility surface area (SASA) at the interface were increased in the complex for coFAM132b, compared to the native complexes and other mutations. ΔG>0 for A136T or P159A mutant indicates that these complexes are unlikely to form in vivo. Interface analysis showed that two hydrogen bonds connecting coFAM132b and insulin B-chain are exactly in the protective hinge at Phe (B24) and Tyr (B26). Of them, the hydrogen bond length between ALA285 and TYR(B26) is only 3.05Å. This strong hydrogen bond contributes to stabilizing the complex (Fig. S5 right). However, these hydrogen bonds collapse completely at the interface between FAM132b and insulin B-chain (Fig. S5 left). Site-specific mutagenesis of FAM132b shows that Ala136 and Pro159 are key sites for insulin-receptor interaction with higher binding constants compared to those of their native counterparts, in which either A136T or P159A mutant displayed no effect on insulin-receptor interaction (Fig. S4C).

## Discussion

We show here that AAV9-mediated expression of FAM132b with A136T and P159A is a safe and effective therapeutic strategy for improving glucose homeostasis and preventing obesity. This is the first demonstration of a therapeutic effect on metabolic disorders in mice with FAM132b gene therapy. A recent study shows that myonectin (FAM132b) deletion promotes adipose lipid storage and reduces hepatic steatosis (35), consistent with our expectation that FAM132b will be taken as a gene therapy target for metabolic disorders.

Skeletal muscle is one of sources producing endogenous FAM132b, which leads to elevated levels of FAM132b in circulation during exercise (27). Due to high-efficiency transduction of the AAV9 serotype for skeletal muscle (36), we used AAV9 vector and muscle-specific promoter to package virus and thus induce the overexpression of FAM132b in skeletal muscle. In general, we observed that skeletal muscle and liver had higher expression of FAM132b with a parallel increase in circulating levels. At the early stage of high-fat diet on 4-week-old mice, the mice firstly exhibited glucose intolerance, followed by insulin resistance and increased EPI sensitivity. This model is closest to the metabolic characteristics of children obesity and insulin resistance. We demonstrated that FAM132b gene therapy in vivo resisted the progression of obesity and metabolic disorders, and altered adrenergic responses in the whole-body, especially in adipose tissues. Contrary to our hypothesis, FAM132b gene therapy also resulted in increased expression of FAM132b in the liver. Gene transfer by the tail vein is likely to transfect AAV9 into multiple organs and tissues, so that the adrenergic response of multiple organs can be modulated together. FAM132b overexpression in liver may also affect hepatic metabolic processes and autophagy (23, 37), and thus regulate the glucose and lipid metabolism of whole-body. The overall effect of FAM132b gene therapy is that reduced adrenergic response of blood glucose, as well as increased adrenergic response of fat, results in an improvement in glucose homeostasis and lipid metabolism.

To reach the therapeutic dose range for AAV9 in vivo, we defined the base titer and set a double dose based on this dose. However, the titer might not reflect the real infectivity of viral particles in vivo, due to batch to batch variations and the type of preparation (38). Double titer transfection did not show double effects of FAM132b overexpression. Indeed, we noticed that the improvement in glucose homeostasis in mice at the higher dose was not significant, but the higher dose of AAV9 was still potential for reducing bodyweight and weight gain. In addition, the lower dose of AAV9: FAM132b did not resulted in a significant increase in FAM132b circulating levels and FAM132b overexpression in skeletal muscle. Although the higher dose of AAV9:FAM132b achieved the transfection effect and altered insulin signaling in fat and skeletal muscle at the protein level, its effects on glucose metabolism was not significant in whole body. In order to minimize the risk of virus transfection, future studies on FAM132b gene transfer in vivo should explore the therapeutic FAM132b RNA levels in target tissues at the lowest titer.

FAM132b has been reported to regulate cellular iron homeostasis, fatty acid transport, fatty acid metabolic process, autophagy and inflammation (25, 26, 37, 39, 40). FAM132b is selected as a potential target for gene therapy because it always ACTS as a metabolic coordinator between tissues and organs. FAM132b (Myonectin) has been identified as a postprandial signal derived from skeletal muscle to integrate metabolic processes in other tissues, and promotes fat transfer from adipocytes to the liver (41). FAM132b-deficient mice exhibited greater fat storage and a striking reduction in liver steatosis (35). Thus, FAM132b gene transfer increases the risk of fatty liver disease by preventing fat accumulation and obesity. The findings of this study further extend our understanding of FAM132b as a metabolic coordinator. In the context of FAM132b gene therapy, the adrenergic response of blood glucose in mice on high-fat diet was decreased, but the adrenergic response of lipolysis was enhanced. These findings indicate that FAM132b not only balances lipid metabolism between different tissues, but also coordinates glucose and lipid metabolism at the whole-body level. We hypothesized that the whole-body effect was related to FAM132b overexpression in the liver, and that targeted injection of AAV9 into skeletal muscle might not result in this effect.

For adipose tissue, increased insulin sensitivity contributes to glucose uptake and utilization, and also contributes to lipogenesis and obesity, so it is of great significance for adipose tissue lipolysis and thermogenesis in adrenergic response. FAM132b promotes fatty acid uptake in adipocytes and hepatocytes and thus improves systemic lipid homeostasis (23). However, if these tissues have lower adrenergic response, it will lead to reduced lipid hydrolysis, fatty liver and obesity. This study showed that FAM132b gene transfer enhanced lipolytic response to EPI in WAT and BAT and increased BAT thermogenesis. These results suggest that FAM132b plays an opposite role in regulating the adrenergic response of blood glucose and adipose tissue. Increasing FAM132b expression leads to lower glycemic response to EPI and higher lipolytic response to EPI in adipose tissues. This regulation indicates a change in fuel selection of the whole-body. We found that gene therapy shifted skeletal muscle fuel selection toward fat oxidation during exercise. The extent of phenotypic improvements induced by FAM132b gene therapy, along with the affinity of AAV9 for skeletal muscle and liver, are promising for the application of FAM132b gene therapy in the clinic for metabolic diseases such as diabetes and obesity. Due to the lack of longer observations in vivo and more delivery routes, preclinical safety studies with different delivery routes need to be carried out for a long time before planning the advancement into human studies.

It is equally important for human to maintain the stability of blood glucose and the sensitivity of blood glucose to EPI. Although studies have shown that hyperglycemia is harmful to multiple organs such as pancreas, kidney, brain, vessels and heart (42-44), the elevated blood glucose under adrenergic stimulation is a necessary response to HPA axis (45, 46). Skeletal muscle is essential for human to escape invasion and capture food in evolution. Changes in gene expression and atypical endocrine of skeletal muscle during strenuous exercise may provide neural feedback and regulate metabolism of other peripheral organs through myokines, and eventually make the best match between sympathetic nervous activity and the metabolic level of peripheral tissues (47-49). Irisin as a myokine produced by skeletal muscle mediates muscle-kidney crosstalk and suppresses metabolic reprograming and fibrogenesis during kidney disease (50). Our study hypothesizes that FAM132b secreted by skeletal muscle and liver enhances adipocytes insulin sensitivity, lipolysis, and thermogenesis through some unknown receptors. Adipose-specific ATGL ablation induced a whole-body shift from lipid metabolism to glucose metabolism by increasing skeletal muscle glucose uptake (51). Thus, adipose tissue lipolysis alters whole-body fuel selection and contributes to glycemic control in adrenergic response. However, AAV9 transfection into adipose tissues was lacking in this study, and FAM132b receptors were not yet known. Structural analysis and ZDOCK results render us predict that ADRB2 and insulin may be a docking site for circulating FAM132b. This docking could be strengthened by codon mutations, thus interfering adrenergic response and the binding of insulin to its receptor. This study suggested that FAM132b gene therapy blunts the glycemic response to EPI, but it may lead to hypoglycemia. This needs to be carefully considered when FAM132b gene therapy is used to treat diabetes.

In conclusion, adeno-associated viral vectors (AAV9) could be used to genetically engineer liver or skeletal muscle to secrete FAM132b. Treatment of mice under high-fat diet feeding with AAV9 encoding FAM132b in vivo resulted in marked reductions in adrenergic response of blood glucose, and enhanced adrenergic response of adipose tissue lipolysis. FAM132b overproduction in healthy mice fed a standard diet promoted skeletal muscle to burn fat during exercise. These therapeutic effects indicate that FAM132b gene transfer with selective codon optimization in vivo might be a valid therapy for obesity that can be extended to other metabolic disorders.

## Materials and Methods

### Animals

C57BL/6J mice were purchased from Shanghai SLAC laboratory Animal Co., Ltd (SLAC, China). Mice were housed and bred in a temperature-controlled room (22∼25°C) on a 12h light/dark cycle in cages. Mice were fed *ad libitum* with a standard diet (MD12031, Medicience diets) or a high-fat diet (45 kcal% fat, MD12032, Medicience diets). Body weight was recorded twice a week. All animal procedures were conducted under protocols approved by the animal experimentation committee of East China Normal University, China.

### Glucose Tolerance, Insulin Sensitivity and EPI Sensitivity Test

Mice were deprived of food for 16 h and then subjected to glucose, insulin and EPI tolerance test (IPGTT, IPITT and IPETT). Blood was collected from a small incision in the tip of the tail (time 0) and then at different time-points after an i.p. injection of glucose (1g/kg), insulin (0.75U/kg), and EPI (0.1 or 0.5 mg/kg). Blood glucose levels were measured using a glucometer (Accu-Check^®^ Active, Roche). Blood lactate and plasma glycerol were determined with enzymatic colorimetric assay following the manufacturers’ protocols (Nanjing Jianchen Biotech).

### Exercise Protocols

Acute exercise was performed on a treadmill for 40 min running. Mice in group were subjected to an exercise program at the speed of 15m/min, which is respectively equal to 80% VO_2_max as measured previously in C57BL/6J mice (52). Immediately after removal from the treadmill running, animals were euthanized by decapitation. Blood was collected and tissues were extracted rapidly and frozen in liquid nitrogen.

### Recombinant AAV vectors

An AAV expression cassette was obtained by cloning, between the ITRs of AAV9, the murine codon-optimized FAM132b coding sequence (rs32077626: A136T; rs30716675: P159A) under the control of the cytomegalovirus (CMV) promoter. A non-coding cassette carrying the CMV promoter but no transgene was used to produce null vectors. Single-stranded AAV9 vectors were produced by transfection in HEK293 cells and purified using an optimized CsCl gradient-based purification protocol that renders vector preps of high purity and devoid of empty capsids.

### Administration of AAV vectors

Recombinant AAV9 encoding FAM132b cDNA sequence under the control of the CMV promoter or FAM132b RNAi sequence were produced in HEK293T cells and purified by a CsCl-based gradient method as described previously (53). A non-coding plasmid was used to produce empty vectors for control mice. For systemic administration, AAV9 vectors were diluted in 200µl of sterilized PBS and injected via the tail vein (∼4.7×10^10^/mouse).

### Histology

Harvested tissues were fixed with 4% paraformaldehyde and then embedded with paraffin. Tissue sections were sliced and stained with haematoxylin and eosin. Slides were digitally scanned using an Aperio ScanScope System to produce high-resolution images. Image J software was used for quantitative analysis of cell size and count.

### Tissues Culture ex vivo

Gastrocnemius muscles, liver tissues, BAT or WAT were extracted from mice and preincubated for 2 h at 30°C in Krebs-Ringer bicarbonate HEPES buffer (KRBH) containing (mM): 129 NaCl, 5 NaHCO_3_, 4.8 KCl, 1.2 KH_2_PO_4_, 1.2 MgSO_4_, 2.5 CaCl_2_, 2.8 glucose, 10 HEPES, and 0.1% BSA at pH 7.4. For ex vivo glucose uptake assay, tissues were incubated for 2 h with insulin 60 mU/mL followed by measurement of 2-deoxy-[^3^H]-glucose uptake as described previously (54). For dose-dependent analysis of glycolysis and lipolysis in tissues ex vivo, tissues were incubated for 15 min with EPI at 0, 1, 5, 10, or 50μg/mL followed by measurement of lactic acid, glycerol or NEFA contents in culture medium. For time-dependent analysis of lipolysis in tissues ex vivo, tissues were incubated for 120-180 min with EPI at 10μg/mL followed by measurement of glycerol or NEFA in culture medium per 30 min (55). Lactate, NEFA and glycerol contents were measured following the manufacturers’ protocols (Nanjing Jianchen Biotech).

### Serum Biochemistry Measurement

Serum samples were gathered from blood collected by cardiac puncture. Serum FAM132b, insulin, leptin, NE, EPI, corticosterone, and ACTH were determined using Mouse ELISA kit (Shanghai Enzyme-linked Biotechnology), respectively. Serum glucose, triglyceride (TG), glycerol were quantified spectrophotometrically using an enzymatic assay (Nanjing Jianchen Biotech) in Infinite 200 PRO Microplate Reader (TECAN).

### Muscle and Liver Glycogen

Gastrocnemius muscle and liver were weighed for less than 100mg, and digested in 30% KOH (m/v) for 15 min at 37°C. Ethanol and Na_2_SO_4_ (75% and 0.32% final) were added to samples to precipitate glycogen overnight at −20°C. Sample pellets were washed twice with 10% KOH and 75% ethanol. Pellets were dissolved with 4 N H_2_SO_4_ at 100°C for 20min and then neutralized with 4 N NaOH. Glucose was determined using Sigma’s glucose assay kit (Nanjing Jianchen Biotech) and was normalized to starting tissue mass.

### Tissue Lipid Metabolites

For glycolytic and citric acid cycle analysis, frozen gastrocnemius samples were pulverized in a liquid nitrogen-chilled mortar. Portions of this frozen tissue were then extracted with 1 mL of a solvent mixture consisting of 8:1:1 HPLC grade methanol:chloroform:water, using a ratio of 1 mL solvent per 30 mg of wet tissue mass. Samples were analyzed by HILIC-TOFMS on an Agilent 7890B as described previously (56). For acylcarnitine analysis, weighed samples were extracted using a mixture of 8:2 of HPLC grade acetonitrile:water, probe sonicated for 20 sec (muscle) and then centrifuged. Supernatants were collected and then dried by vacuum centrifugation at 45°C. Dried samples were reconstituted in a 9:1 mixture of mobile phases A/B used for the acyl-carnitine analysis as described previously (56).

### Quantitative PCR

Total RNA was isolated from tissues using TRIzol (Invitrogen), according to the manufacturer’s instructions. RNA was reverse-transcribed (cDNA Synthesis Kit, TOYOBO, Osaka, Japan). cDNA was amplified by real-time PCR in a total reaction volume of 20μl using SYBR Green Realtime PCR Master Mix (QPK-201; TOYOBO, Osaka, Japan). Real-time PCR reactions were cycled in StepOne™ Real-Time PCR System (Applied Biosystems) using the following primers in Supplemental Materials. Target gene expression was normalized to housekeeping gene 18S or GAPDH and expressed as 2^−ΔΔct^ relative to the control group, respectively.

### Western blot analysis

For frozen tissue, lysates were prepared using a RIPA lysis buffer (50mM Tris (pH7.4), 150mM NaCl, 2mM EDTA, 1% Nonidet P-40, 50mM NaF, 0.1% SDS and 0.5% sodium deoxycholate with PhosStop Phosphatase-Inhibitor Cocktail tablet and protease-inhibitor cocktail tablet (Roche). The homogenate was centrifuged at 4°C for 10 min at 14,000g and the supernatant was used for western blot analysis. The protein content of the supernatant was quantified using bicinchoninic acid reagents and BSA standards. Equal amounts of protein were separated using a polyacrylamide SDS-PAGE gel. After SDS-PAGE, proteins were transferred to PVDF membrane. The membrane was blocked for 1 h at room temperature followed by incubation overnight at 4°C with primary antibodies including FAM132b (AVISCERA BIOSCIENCE), PI3K, PTEN, p70S6K, Akt, p-Akt (Ser473), UCP1 (Cell Signaling, USA), beta-Actin or GAPDH (Servicebio, China). After overnight incubation, blots were incubated with HRP-conjugated secondary antibodies at a dilution of 1:5,000 for 1 h at room temperature. Bands were visualized by ECL plus according to the manufacturer’s instructions (Thermo Scientific) and quantified using Image J software.

### Modeling the FAM132b structure

The amino acid sequence for FAM132b was obtained from UniProt database in FASTA format to be utilized for the prediction of FAM132b structure. SOMPA was used to predict the secondary structure of FAM132b and codon-optimized FAM132b (coFAM132b) (57). In this study, only the 25–340 part of the sequence was used in homology modeling. Since SWISS-MODEL has less than 30% sequence identity (58), we next used I-TASSER server to model the unknown structure of FAM132b (59-61). coFAM132b was modeled in the same way. The quality of the selected model was further validated using PROCHECK (62), a tool that generates a Ramachandran plot and discusses the stereochemical aspects of the FAM132b. VMD 1.9.3 was used to visualize FAM132b structure and analyze hydrogen bonds (63).

### Determining the binding modes of FAM132b to Target using ZDOCK

To determine the probable binding of FAM132b and its potential targets, we used ZDOCK (64). As initial structures, the FAM132b structure obtained by I-TASSER was used. ZDOCK performed a global grid search in position and orientation of proteins with respect to FAM132b and its mutants, with local docking to explore the neighborhood of target with the lowest energy in detail. The ZDOCK server is set to produce 500 structures of complexes in global docking and a smaller subset in restricted docking. The rank first reasonable structure of the FAM132b-target complexes is analyzed by PDBePISA (65). To verify the importance of selected amino-acid residues in the obtained most probable model of the FAM132b-target complexes, selected residues of FAM132b were replaced with A136T and P159A, and the stability of the resulting complexes was assessed by PDBePISA. VMD 1.9.3 was used to visualize the docking model and analyze hydrogen bonds at the interface (63).

### Quantification and Statistical Analysis

Data are presented as means±SEM. Significance was assessed by Student’s t-tests and/or ANOVA with Tukey post hoc tests for multiple comparisons where appropriate. Statistical analysis were performed with GraphPad Prism 8.1, and differences were considered significant when p<0.05. Significances are indicated in the figures according to the following: *p < 0.05, **p < 0.01, and ***p < 0.001. Statistical parameters can be found in the figure legends.

## ACKNOWLEDGMENTS

We thank Dr. Li Rui, the director engineer of Genomeditech (shanghai) Co.,LTD, for designing and producing recombinant AAV vectors of serotype 9 encoding FAM132b cDNA sequence or RNAi sequence. We thank Dr. Wu J.L. and Dr. Liu Y.M., technician of the Instrumental Analysis Center of Shanghai Jiaotong University, for their expertise and conversations related to body composition analyses and metabolic assays. Supported by Natural Science Foundation of China grants 31871208 and 31300977, Shanghai Natural Science Foundation (18ZR1412000), the Fundamental Research Funds for the Central Universities (40500-20104-222288) and the Key Laboratory Construction Project of Adolescent Health Assessment and Exercise Intervention of Ministry of Education, China (No. 40500-541235-14203/004).

## AUTHOR CONTRIBUTIONS

Z.Q. and W.L. conceived, designed, performed, and interpreted experiments and made figures. J.X., X.X., J.L., W.L., L.L., Z.C., and Z.H. performed animal care, treadmill exercise, glucose tolerance tests, insulin tolerance tests, EPI tolerance tests, real-time PCR, and data collection, interpreted a part of experiments and made some figures. X.Z., Q.Z., X.L., L.C., B.J. performed a part of animal care, acute exercise, metabolites analysis, and a part of data processing. Y.Z., J.L. performed structural analysis and ZDOCK. S.D. participated in interpreting results and supervising the experimental plan. Z.Q. and W.L. wrote and revised the manuscript.

